# Inducing aggresome and stable tau aggregation in Neuro2a cells with an optogenetic tool

**DOI:** 10.1101/2024.05.07.592868

**Authors:** Shigeo Sakuragi, Tomoya Uchida, Naoki Kato, Boxiao Zhao, Toshiki Takahashi, Akito Hattori, Yoshihiro Sakata, Yoshiyuki Soeda, Akihiko Takashima, Hideaki Yoshimura, Gen Matsumoto, Hiroko Bannai

## Abstract

Tauopathy is a spectrum of diseases characterized by fibrillary tau aggregate formation in neurons and glial cells in the brain. Tau aggregation originates in the brainstem and entorhinal cortex and then spreads throughout the brain in Alzheimer’s disease (AD), which is the most prevalent type of tauopathy. Understanding the mechanism by which locally developed tau pathology propagates throughout the brain is crucial for comprehending AD pathogenesis. Therefore, a novel model of tau pathology that artificially induces tau aggregation in targeted cells at specific times is essential. This study describes a novel optogenetic module, OptoTau, which is a human tau with the P301L mutation fused with a photosensitive protein CRY2olig, inducing various forms of tau according to the temporal pattern of blue light illumination pattern. Continuous blue light illumination for 12 h to Neuro2a cells that stably express OptoTau (OptoTauKI cells) formed clusters along microtubules, many of which eventually accumulated in aggresomes.

Conversely, methanol-resistant tau aggregation was formed when alternating light exposure and darkness in 30-min cycles for 8 sets per day were repeated over 8 days. Methanol-resistant tau was induced more rapidly by repeating 5-min illumination followed by 25-min darkness over 24 h. These results indicate that OptoTau induced various tau aggregation stages based on the temporal pattern of blue light exposure. Thus, this technique exhibits potential as a novel approach to developing specific tau aggregation in targeted cells at desired time points.

**Significance:** This study developed an approach to manipulate tau aggregation in a blue light-dependent manner using cells that stably express OptoTau, which is an optogenetic tool based on the CRY2olig module. Tau accumulation in aggresomes or stable tau aggregation were selectively induced by blue light illumination conditions. These results are crucial as they provide a new technological basis for establishing a singular point of tau aggregation in specific targeted cells at a particular time.

## Introduction

Tauopathy is a spectrum of diseases that are characterized by neurofibrillary tangles (NFTs), which are filamentous tau protein aggregates that are formed in neurons and glial cells in the brain [1]. Tauopathy manifests as various disorders, such as Alzheimer’s disease (AD), Pick’s disease, progressive supranuclear palsy (PSP), corticobasal degeneration (CBD), and chronic traumatic encephalopathy (CTE), despite being caused by abnormalities in the same molecule tau. Microtubule-binding protein tau in healthy neurons exerts physiological roles, such as neurite outgrowth and axonal transport regulation by motor proteins. However, tau detaches from microtubules and forms intracellular tau aggregates when undergoing post-transcriptional modifications, including excessive phosphorylation [2]. The hyperphosphorylation of tau as a trigger of NFT is initially formed in the neurons in locus coeruleus (LC) and entorhinal cortex (EC) in AD, which is the most prevalent type of tauopathy [3,4]. This tau aggregation spreads with age into the transentorhinal region, hippocampus, and cerebral cortex (temporal lobe, prefrontal cortex, and higher sensory association neocortex) [4]. Tau aggregation progression is closely related to the clinical pathology development of AD [5,6].

Numerous model experimental systems have helped understand the association between degenerative tau accumulation and higher brain functions in AD and the mechanisms underlying tau pathology expansion. Tau aggregate formation, synapse loss, neurodegeneration, and brain atrophy have been observed in a few months in mice that overexpress neurons with human tau with 301st proline to serine or leucine (P301S or P301L) mutations found in familial frontotemporal dementia [7-10]. These phenotypes exhibit common pathologies in human patients; thus, these mice were selected as an AD model, which help elucidate the mechanism of toxicity from degenerated tau. Tau filaments polymerized *in vitro* and insoluble tau fractions from patients with AD injected into human tau-overexpressing mice form hyperphosphorylated tau and tau aggregates in distant brain regions with neuronal connections [11-13]. Additionally, tau extracted from patient brains induces pathology, even in non-transgenic mice [14,15]. The spread of degenerated tau to healthy neurons has occurred through synaptic contact or by another cell uptake of degenerated tau that is extracellularly released depending on a synaptic activity [16, 17]. These experiments have helped elucidate some aspects of tauopathy pathogenesis as model systems that rationally reproduce some of the phenotypes observed in AD and other human tauopathies.

Recently, Nagai et al. defined a “singularity phenomenon” as a biological phenomenon in which a small number of rare cells, named “singularity cells,” significantly change a complex system [18,19]. Therefore, neurons that first accumulate hyperphosphorylated tau and NFTs in LCs and ECs were considered “singularity cells” in AD [20]. Addressing the following questions is considered crucial for understanding the onset mechanisms of AD: “How do these “singularity cells,” which initially accumulate tau, arise?” and “How is tau degeneration transmitted from the “singularity cells” to the entire cortex?” However, the current experimental system of tauopathy models faces several challenges in addressing this issue. Although the brain regions that tend to aggregate tau have been identified in the tauopathy mice model with human tau overexpression, why and how the first “singularity cells” start tau aggregation in that brain region still remains unclear. More seriously, the loss of endogenous gene was detected in current animal models overexpressing human tau, and it should be reconsidered whether tau abnormalities are the only cause of pathological phenotypes in these animals [10]. Another AD model, obtained by exogenous tau seed injection [14] [15], bypasses the initial process of AD pathogenesis. Tau seed uptake is entirely random, and determining the cell from which the pathology originates is impossible. A new model system is required to establish a singular point of tau aggregation at a targeted time in specifically targeted cells to elucidate the tau pathology process from the singularity cells.

This study employed optogenetics to induce tau aggregation in targeted cells. CRY2, which is a photoreceptor in *Arabidopsis*, and its derivative CRY2olig are optogenetic modules that reversibly form oligomers during blue light exposure [21]. Recently, researchers have introduced CRY2 to trigger protein aggregation in neurodegenerative diseases, including tau [22-25]. CRY2 and tau fusion proteins accumulated on microtubules by light-induced liquid-liquid phase separation (LLPS) [23]. Blue light illumination forms tau droplets and oligomers in primary cortical culture neurons of mice that overexpress tau fused with CRY2olig, which is a higher capacity for homo-oligomer formation [24]. Recently, we have shown that the formation of LLPS droplets and tau oligomers was induced by blue-light illumination for 24 h in Neuro2a cells overexpressing “OptoTau,” a fusion of human tau with the P301L mutation and CRY2olig [25,26]. The present study manipulated intracellular tau by blue light illumination in Neuro2a cells that stably express OptoTau. OptoTau, which is inserted into the safe-harbor region by CRISR/Cas9 technique, underwent various blue light illumination temporal patterns, including eight-day illumination. Here, we report that different blue light illumination patterns enable the same optogenetic tool, OptoTau, to cause different responses, including accumulation of tau in the aggresome and creation of stable tau aggregates.

## Materials and Methods

### OptoTau and CRY2olig SNAP-tag knock-in into Neuro2a cells

OptoTau involves the optogenetic module CRY2olig [21], a SNAP-tag, and a full-length human tau (2N4R) with a P301L (Figure 1A). ROSA26 MCS was utilized to establish the OptoTau-donor vector, OptoTau gene under the cytomegalovirus promoter, and Green Fluorescent Protein (GFP) and puromycin-resistant genes under the EF1α promoter. Lipofectamine 3000 (Life Technologies) was used to transfect the OptoTau-donor and mROSA26sgRNA/Cas9 vectors (GeneCopoeia), such as the guide RNA and Cas9 sequences, into mouse neuroblastoma-derived Neuro2a cells (JCRB Cell Bank). Positive selection was performed by adding 400 μg/ml G418 (Nacalai tesque) on day one post-transfection and 1 μg/ml puromycin (Nacalai tesque) after 4 days. GFP was utilized as a gene insertion marker, and GFP-positive cells were isolated into a 96-well plate. Single GFP-positive cell colonies were replated into a 24-well plate and further spread on a 10-cm culture dish. This study used one clone as OptoTau knock-in (OptoTauKI) cells after confirming OptoTau expression.

**Figure 1.**
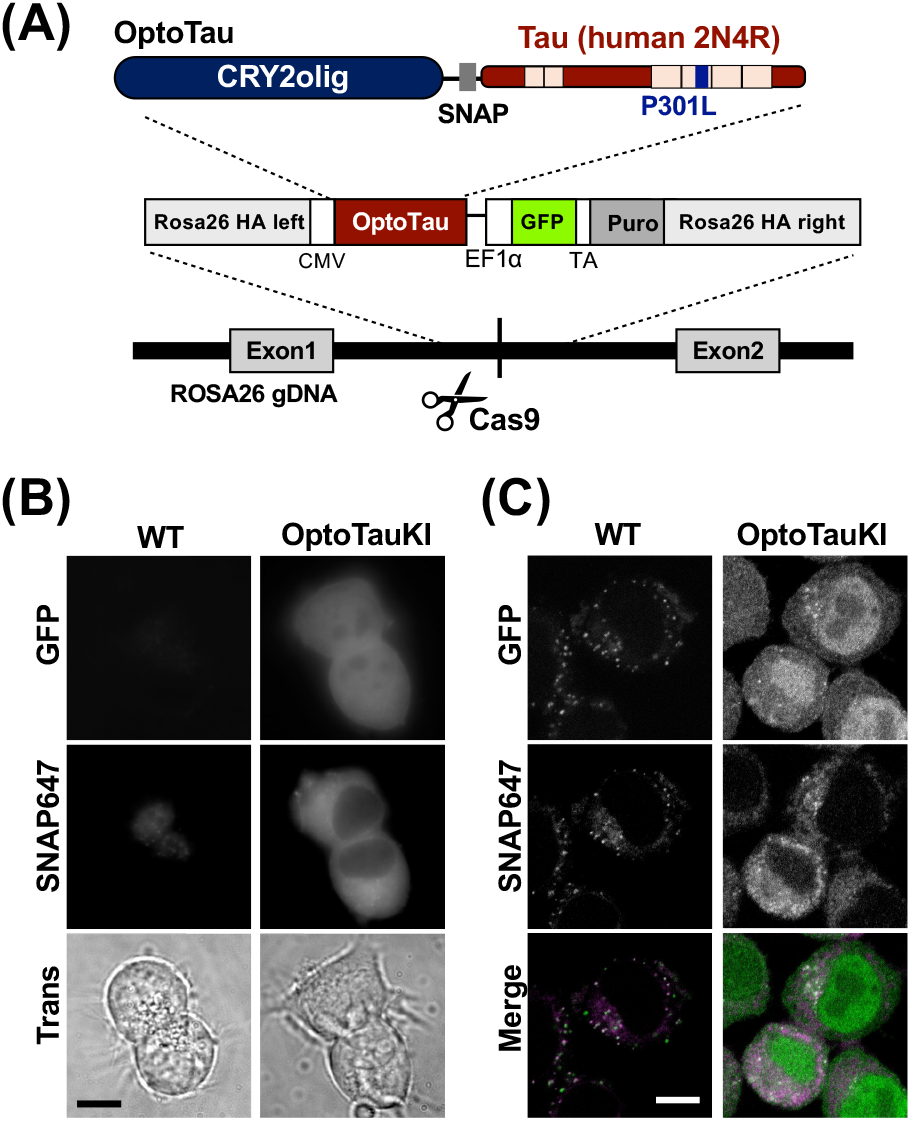
OptoTau knock-in Neuro2a cells. **(A)** Top: Design of OptoTau, consisting of CRY2olig, SNAP-tag, and human2N4R tau with P301L mutation; bottom: conceptual diagram of OptoTau knock-in to mouse ROSA26 locus with the CRISPR/Cas9 system. The donor vector contains OptoTau expressed under the CMV promoter and GFP and puromycin (Puro) sequences for screening. **(B)** Live cell imaging of GFP and SNAP647 fluorescence in Neuro2A cells (WT) and OptoTau knock-in Neuro2a cells (OptoTauKI). The SNAP647 signal represents OptoTau localization. Transmission images (Trans) were obtained from the same field of view. Bar indicates 10 μm. **(C)** Confocal images of GFP and SNAP647 fluorescence in Neuro2A cells (WT) and Neuro2a cells knocking in OptoTau knock-in (OptoTauKI). Cells were fixed with 4% PFA. Bar indicates 10 μm.

As a control, CRY2olig SNAP-tag, the protein with tau sequence removed from OptoTau, was knocked-in to the ROSA26 locus in Neuro2a cells following the above procedure, except for single-colony isolation. Positive selection was performed by adding 1 or 2 μg/ml puromycin for a week, starting 7 days post-transfection.

### Cell maintenance

OptoTauKI and Neuro2a wild type (WT) were maintained in Dulbecco’s Modified Eagle’s medium (4.5 g/l of glucose; Nacalai tesque) supplemented with 10% fetal bovine serum, penicillin (100 units/ml), and streptomycin (100 μg/ml) mixture, in 5% CO_2_ at 37 °C environments. OptoTauKI cells were kept under light-shielded conditions to prevent irregular CRY2olig conformational changes caused by room light. Cells were plated on a 35-mm glass bottom dish (MATSUNAMI) coated with 0.04% polyethyleneimine (PEI, Sigma–Aldrich) for the live imaging experiment. Cells were plated on an 18-mm cover glass precoated with 0.04% PEI for fixation. The density of light-exposed cells was maintained at 60%–80% confluence, except in the experiment where the cells were incubated for 48–72 h after light illumination.

### Blue light illumination

A blue light illumination device (TH2-211×200BL, CCS Inc.) was set in a 5% CO2 incubator at 37°C. Cell plates were placed on a transparent plastic box with a 30-mm height on the light-emitting device. The light intensity was 0.24 ± 0.04 mW/cm^2^ throughout all experiments. This study conducted three illumination protocols, (1) single 12-h illumination (single 12 h), (2) 16-h break followed by 8 cycles of 30-min illumination of 30-min light off per day for up to 8 days (break and repeat), and (3) 48 cycles of 5-min illumination at 25-min light off per day (48 repeats). An automatic timer controlled the illumination timing.

### Labeling of OptoTau using SNAP-tag

OptoTau molecules were fluorescently labeled by covalently binding a fluorescent-labeled SNAP ligand to a SNAP-tag inserted into the OptoTau. SNAP-Cell 647-SiR (New England BioLabs) of 0.1 μM was administered into the culture medium, and OptoTauKI or WT was employed for 12–18 h. Cells were washed with medium and then utilized for live cell imaging or immunocytochemistry.

### Immunocytochemistry

Primary antibodies involved anti-total tau antibody (RTM38), an antibody recognizing the initial structural change of pathological tau (MC1, gifted from Prof. Peter Davies [Albert Einstein College of Medicine Medicine]), anti-phosphorylated tau antibody (AT8), anti-αTubulin antibody, anti-γTubulin antibody, anti-HDAC6 antibody, and anti-Dynein antibody that were obtained and used at the dilution indicated in Supplemental Table 1. Phosphate buffered saline (PBS) was used for dilution and washing throughout the immunocytochemistry. Cells were generally fixed with 4% paraformaldehyde (PFA) for 15 min at room temperature (RT: ∼24°C) and permeabilized with 0.1% Triton X-100 for 3 min at RT. For microtubule or aggresomal marker immunostaining, cells were fixed with ice-cold acetone/methanol (acetone: methanol = 1:1) for 3 min at −20°C, followed by post-fixation with 4% PFA (15 min, RT). To remove soluble tau in the cytosol, cells were fixed with ice-cold methanol for 20 min at −20°C [27]. Cells were incubated with 5% bovine serum albumin (BSA) for 30 min at RT for blocking after fixation and permeabilization, followed by primary antibody treatment in 2.5% BSA for 2 h for AT8 antibody or 1 h for other antibodies at RT. Cells were incubated with secondary antibodies (5 μg/ml in 2.5% BSA) for 40 min at RT after washing with PBS. Secondary antibodies included anti-rat IgG (H + L) Alexa Fluor® 555 conjugate, anti-rat IgG (H + L) Alexa Fluor® 647 conjugate (Cell signaling) or goat anti-Rat IgG (H + L) cross-adsorbed (Alexa Fluor™ 594), goat anti-mouse IgG (H + L) cross-adsorbed (Alexa Fluor™ 647), and goat anti-mouse IgG (H + L) cross-adsorbed (Alexa Fluor™ 555) (Thermo Fisher Scientific). VECTASHIELD mounting medium (Vector Laboratories) was used to mount coverslips.

### Observation by epifluorescence and confocal laser scanning microscopies

An inverted microscope (IX73; Evident) equipped with a high-intensity LED driver (DC4104, Thorlabs) and a digital CMOS camera (ORCA-Fish 4.0; HAMAMATSU) was utilized to observe immunocytochemically or SNAP-tagged cells, set at 60× oil immersion objective (NA 1.42; Evident), 1.6× optical zoom, and 5–10 fields of view captured from a single cover glass. Confocal laser scanning microscope (FV3000; Evident) images were obtained with an oil immersion objective (×60, NA1.42; Evident) at 5× digital zoom.

### Western blotting

A sample buffer solution of 250μl with 2-Mercaptoethanol (30566-22, Nacalai Tesque) was directly added to WT Neuro2a or OptoTau KI cells plated in 3.5 cm and lysed by sonication at room temperature immediately after completing blue light illumination. Samples were loaded onto a 5%–20% gradient gel, electrophoresed at 20 mA, and transferred to a polyvinylidene fluoride membrane (Bio-Rad). The membrane was blocked with 3% BSA with 0.05% Tween-20 and reacted with an anti-tau antibody RTM38, anti-phosphorylated tau antibody AT8, and anti-β-actin antibody (Supplemental Table 1), at RT for 1 h (or 4°C O/N) after transfer. The membrane was incubated with the secondary antibodies conjugated with peroxidase (1:10,000, Jackson Immuno Research) at RT for 1 h after washing three times with PBST (0.05% Tween20 in PBS). A luminol-based chemiluminescence assay kit (Chemi-Lumi One L, Nacalai tesque) and chemiluminescence detection system (ChemiDoc Touch MP imaging system, Bio-Rad) were utilized to detect chemiluminescence signals.

### Quantitative analysis of the percentage of cells with aggregations and spatial distribution of tau aggregation

Quantitative image analysis was performed using MetaMorph (Molecular Devices) or ImageJ (NIH) image analysis software. The percentage of cells with stable tau aggregates in the field of view was calculated per image of PFA-fixed cells, which were subjected to illumination of 8 cycles of 30-min illumination and 30-min light off per day for 6 or 8 days. The cells with the aggregates were defined as those with a maximum RTM38 intensity larger than “the average RTM38 intensity + 3 standard deviations” of cells in the dark condition.

The relative phosphorylation level, which indicates the phosphorylation level of tau in the cell, was defined as the ratio of the average fluorescence intensity of AT8 and RTM38, normalized by the median AT8/RTM38 value in the dark condition cells of each batch.

The difference in the spatial distribution of tau aggregation between cells with single 12-h continuous blue light illumination and cells with repetitive 5-min illumination with 25-min intervals was analyzed using the size of tau aggregation and Gini coefficient, an inequality indicator [28]. This analysis utilized tau images labeled with RTM38 antibody at the focal plane where the nucleus is most clearly visible.

The size of tau aggregates was calculated as the width of the brightest aggregation divided by the width of the nucleus, both of which were measured along the straight line through the center of the nucleus and the brightest point in the aggregation (Supplemental Figure 1A).

We calculated the Gini coefficient from the Lorentz curve plotted from the pixel fluorescence intensity distribution based on previous research to quantify the inequality of tau aggregation fluorescence intensity [29]. The entire cell was selected as a region of interest (ROI), and the fluorescence intensity distribution of the pixels within the ROI was determined (Supplemental Figure 1B). The average nuclear fluorescence intensity (μNuc) was measured, and pixels with fluorescence intensities of ≥2 μ_Nuc_ were defined as tau-positive signals and were subsequently subjected to analysis. A new histogram of the number of pixels, where each bin has a width of 0.1 μ_Nuc_, was established from the extracted pixels (Supplemental Figure 1C). This analysis redefined the fluorescence intensity of the i-th bin as i. Pixels with fluorescence intensity of >10 times that of μ_Nuc_ (10 μ_Nuc_) were included in the bin of the 10-μ_Nuc_ category and the total number of bins was set to 81. The total number of pixels *N* and the total amount of fluorescence *T* can be expressed as 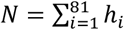, 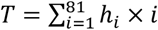, respectively, for this histogram, if the number of pixels in the i-th bin is *hi*. The Lorentzian curve *L(pj)*, which plots the normalized fluorescence intensity against the normalized number of pixels, was expressed as follows (Supplemental Figure 1D):

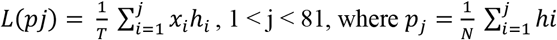

The Gini coefficient was calculated as twice the area (S, gray) between the Lorenz curve (solid line) and the perfect equity curve (blue dotted line).

### Statistics

Figure legends indicate the numbers of data. The Wilcoxon signed-rank test was used to compare Western blots with and without blue light illumination and the Mann–Whitney U-test was utilized for the percentage of cells with stable tau aggregates. Kolmogorov–Smirnov test was conducted to compare the percentage of tau aggregates in the perinuclear region for each light exposure pattern. The *p*-values in each figure are NS: **: *p* < 0.01, ***: *p* < 0.001, ****: *p* < 0.0001.

## Results

### Establishment of OptoTau knock-in (OptoTauKI) cells

OptoTau is a protein fused to CRY2olig [21], which forms multimeric complexes upon blue light exposure, with human 2N4R tau carrying the P301L mutation (Figure 1A). We have previously demonstrated that transient OptoTau overexpression into Neuro2a and blue light exposure induced tau clustering [25] [26]. This study established Neuro2a cells that stably express OptoTau and investigated the effects of blue light illumination over a longer period than in previous studies. The CRISPR/Cas9 system was utilized to introduce OptoTau into the mouse Safe Harbor ROSA26 region. We developed the cells to express OptoTau under the CMV promoter and GFP for screening under the EF1α promoter (Figure 1A). We isolated GFP-positive single colonies and defined them as OptoTau knock-in (OptoTauKI) cells. We conducted fluorescent labeling of SNAP-tag incorporated into OptoTau to confirm OptoTau expression and localization. Both GFP and SNAP647 labeling exhibited no signal in non-genome-edited Neuro2a cells (WT), whereas GFP was observed throughout the OptoTauKI cells, and SNAP-tag signal was found in the cytoplasm (Figure 1B). Confocal imaging indicated localized OptoTau in the cytoplasm, which avoid the nucleus (Figure 1C). These results reveal that OptoTau is inserted in OptoTauKI cells and the expressed OptoTau resides mainly in the cytoplasm.

### Filament-like cluster formation of tau by 12-h blue light exposure

We then investigated the behavior of OptoTau in OptoTauKI cells under blue light illumination. OptoTauKI cells were continuously exposed to blue light for 12 h with a blue LED in a CO_2_ incubator (single 12-h illumination condition), and then tau localization was examined by SNAP staining or immunofluorescence staining. Both the SNAP signal and the signal of total tau visualized by the RTM38 antibody exhibit uniform distribution in the cytoplasm in OptoTauKI cells cultured in the dark (Figure 2A, Dark). In contrast, OptoTau labeled with SNAP647 in OptoTauKI cells exposed to blue light for 12 h formed thick filament-like structures, also recognized by RTM38 antibody (Figure 2A, single 12 h). Single-molecule localization microscopy has demonstrated that tau forms oligomers on microtubules [30]. We conducted multicolor immunofluorescence staining of tau and microtubules in the same cells to identify whether the filamentous tau formed by blue light illumination co-localizes with microtubules. OptoTauKI cells illuminated with blue light for 12 h were fixed with acetone methanol, and microtubules were labeled with α-tubulin antibody and tau with RTM38 antibody (Figure 2B). The filamentous tau signal largely co-localized with that of α-tubulin (Figure 2B, Merge).

**Figure 2.**
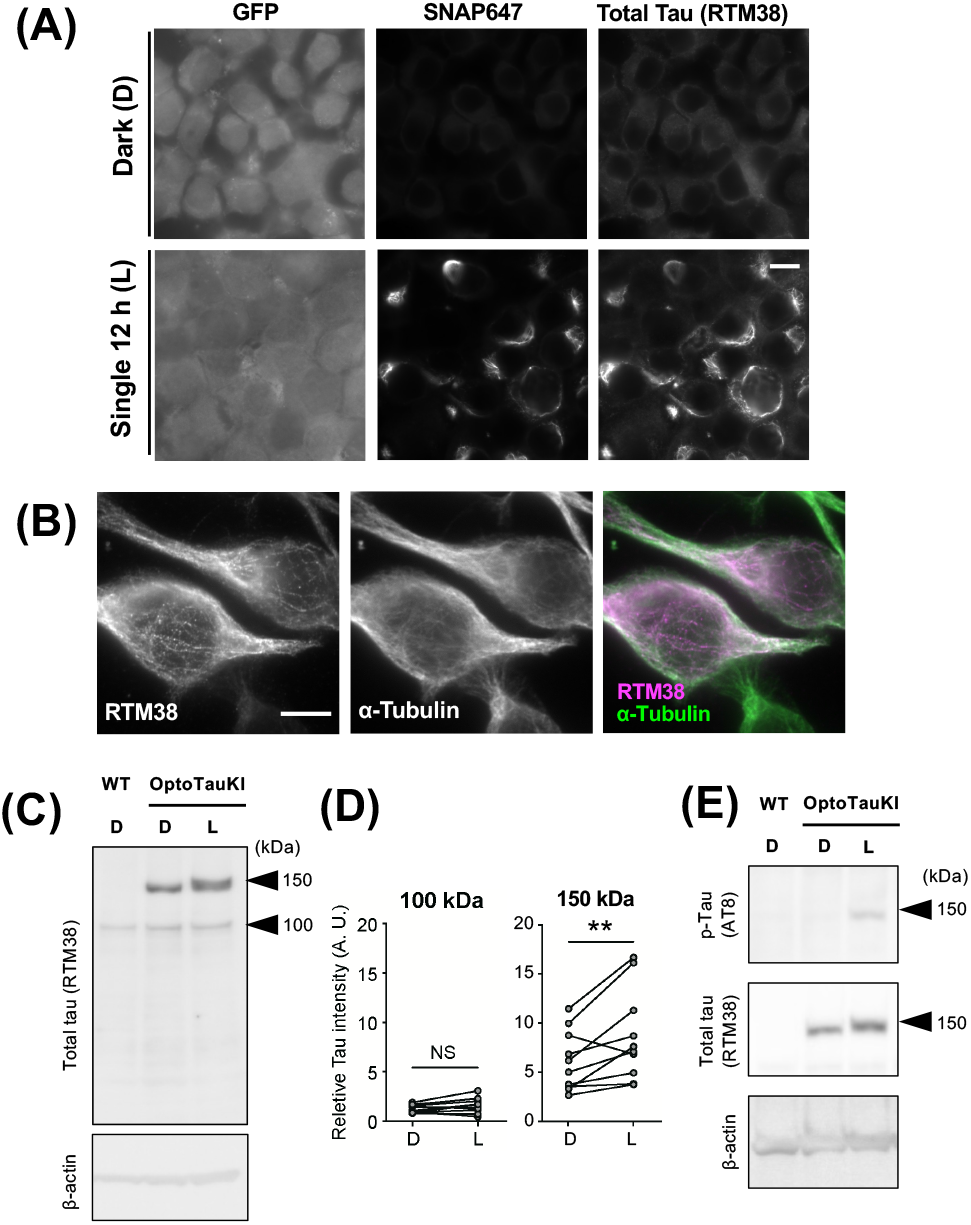
Effect of 12-h blue light illumination on Tau in OptoTau knock-in cells. **(A)** Subcellular localization of GFP (left), OptoTau (middle, SNAP647), and total tau shown by immunofluorescence staining with RTM38 antibody in OptoTau knock-in cells. Top: cells kept in the dark condition (Dark [D]); bottom: cells fixed immediately after a single 12-h blue light illumination (Single 12 h [L]). Scale bar indicates 10 μm. **(B)** Immunofluorescence staining images of microtubules (middle, α-tubulin) and total tau (left, RTM38) in OptoTauKI cells immediately after a single 12-h blue light illumination. The right panel exhibits the merge of these signals (a-tubulin [green], total Tau [magenta]). Scale bar indicates 10 μm. **(C)** Western blotting analysis of Tau using RTM38 antibody in Neuro2a cells (WT) and OptoTauKI cells (OptoTauKI). D: Cells under dark conditions; L: cells with a single 12-h illumination. Bands were detected at 150 kDa and 100 kDa. β-actin was used as a loading control. **(D)** Quantification of 100 kDa (left) and 150 kDa (right) tau. The ratio of β-actin-normalized tau band intensity to that of Neuro2a cells (WT) was defined

Furthermore, a Western blot with RTM38 antibody under reducing conditions was conducted to confirm the tau expression level in OptoTauKI cells. Only a 100-kDa band was detected in Neuro2a cells (WT), whereas two bands of 100 kDa and 150 kDa were detected in OptoTauKI cells (OptoTau) with or without a single 12-h blue light exposure. This was supported by Western blotting using another tau antibody, 5B10, which recognizes different epitopes from RTM38, detecting a large 150-kDa band, a smear band of approximately 100 kDa, and a smear band of 50 kDa (Supplemental Figure 2). These results indicate that the 150 kDa band represented the knocked-in OptoTau and that the 100-kDa band recognized by RTM38 was a protein with a tau epitope. The intensity of the 100-kDa and 150-kDa bands was quantified to investigate the changes in the amount of OptoTau protein with or without blue light illumination. Band intensities were normalized by β-actin, a loading control, and calculated as “relative tau intensity” relative to the value of WT. The relative tau intensity of the 100-kDa band was not different between the cells with and without a single 12-h blue light illumination (Figure 2D, 100 kDa), whereas the relative tau intensity of the 150-kDa band was higher in the blue light group in 9 out of the 10 samples (Figure 2D, 150 kDa). These results reveal the increased amount of OptoTau protein in OptoTauKI cells by the 12-h blue light illumination. The apparent molecular weight of the 150-kDa band corresponding to OptoTau increased after blue light illumination (Figure 2C, Supplemental Figure 2), indicating the possible post-translational modification of tau protein. Indeed, the AT8 antibody, which recognizes pathological phosphorylation, detected the 150-kDa band only under the light condition (Figure 2E). This result indicates that blue light illumination causes pathological phosphorylation of OptoTau-knocked into Neuro2a cells and is consistent with previous findings in the cells overexpressing OptoTau via transfection [25].

### Tau dynamics in OptoTauKI cells after blue light illumination termination

A single 12-h blue light irradiation induced filament-like structures in OptoTauKI cells (Figures 2A and B). Concurrently, Western blot detected no tau aggregates with large molecular weights, which could be oligomers (Figure 2C). Cells were fixed at 0, 24, 48, and 72 h after blue light termination to investigate the stability of filamentous tau formed after the 12-h blue light illumination. The subcellular localization of tau and microtubules was examined with immunofluorescence staining (Figure 3A). Filament-like tau co-localized with microtubules in OptoTauKI cells immediately after blue light termination, exhibiting a strong signal in their part (Figure 3B, 0 h, arrowheads). A bright spot with dense tau was formed in the vicinity of the nucleus 24 h after blue light termination (Figure 3B, 24 h, arrows). Bright spots remained at 48 and 72 h, but the tau bright spot was fragmented in most cells. These results reveal that the filamentous tau formed in OptoTauKI cells after a single 12-h illumination is an unstable structure that dynamically changes morphology over time.

**Figure 3.**
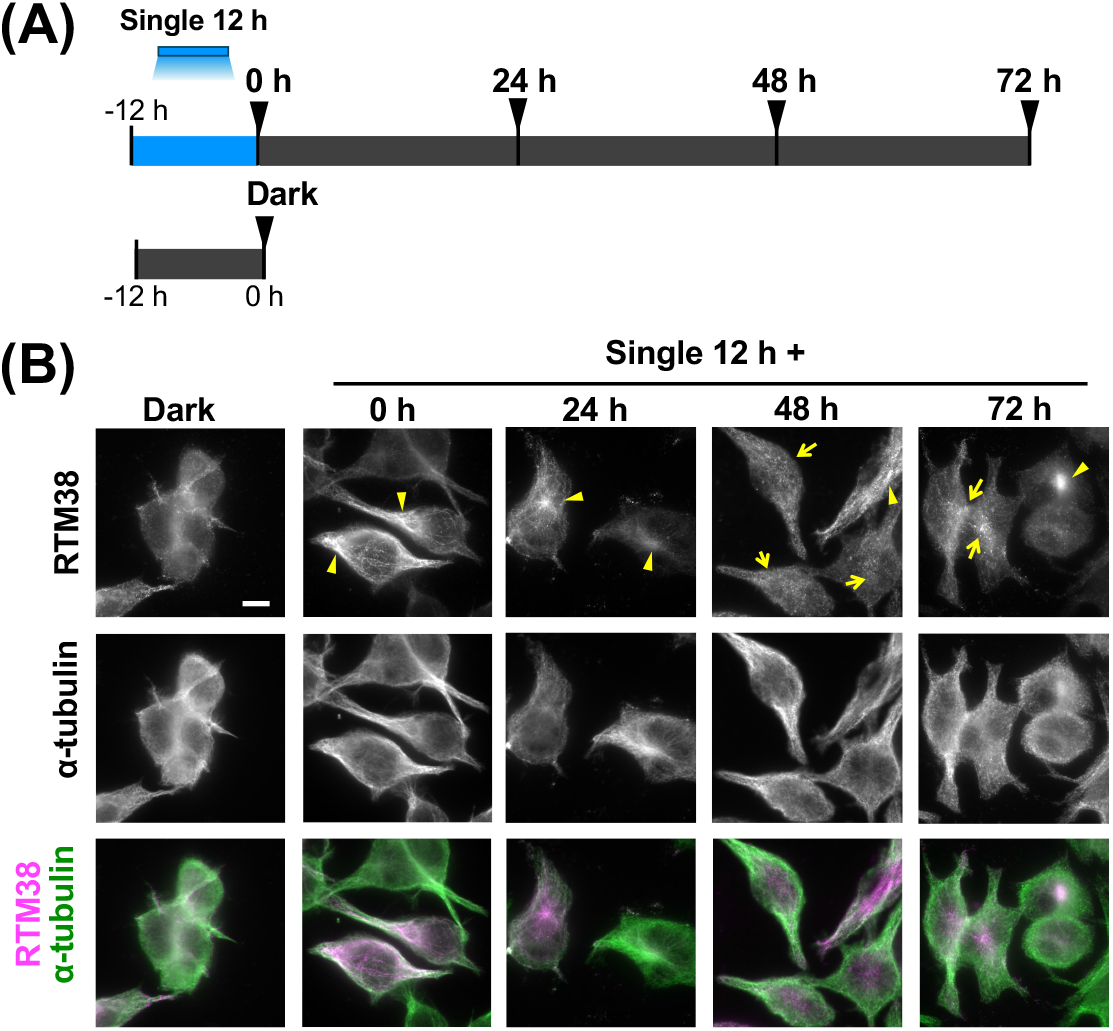
Localization change of Tau after blue light illumination termination. **(A)** Schematic illustration of the experimental design. Top: localization of total tau in OptoTauKI cells cultured in the dark after a single 12-h blue light illumination verified by immunofluorescence staining with the RTM38 antibody; arrowhead represents the timing of cell fixation. Dark indicates cells cultured in the dark for 12 h. (B) Immunofluorescence images of total tau and microtubules immediately (0 h), 24 h, 48 h, and 72 h after blue light exposure termination. Top: RTM38 antibody, middle: α-tubulin antibody, bottom: merge image. Yellow arrowheads indicate large bright spots of tau; arrows indicate fragmented bright spots of tau. Scale bar indicates 10 μm.

### Co-localization of tau with aggresome markers after blue light illumination

Proteasome inhibition and uptake of tau-paired helical protofilaments from patients with AD have induced aggresomes, which are inclusion bodies for misfolded proteins [31] [32]. Aggresomes are formed in the microtubule organizing center (MTOC) located in the vicinity of the nucleus [33]. We conducted immunofluorescence staining for γ-tubulin in the MTOC, together with tau, to investigate the possibility that the perinuclear tau accumulation in OptoTauKI cells after a single 12-h blue light exposure is an aggresome. The tau accumulation in the vicinity of the nucleus observed immediately (0 h) and 24 h (24 h) after blue light termination was co-localized with γ-tubulin in MTOCs (Figure 4A, arrowheads). This result reveals the MTOC as the site of tau accumulation after blue light termination.

**Figure 4.**
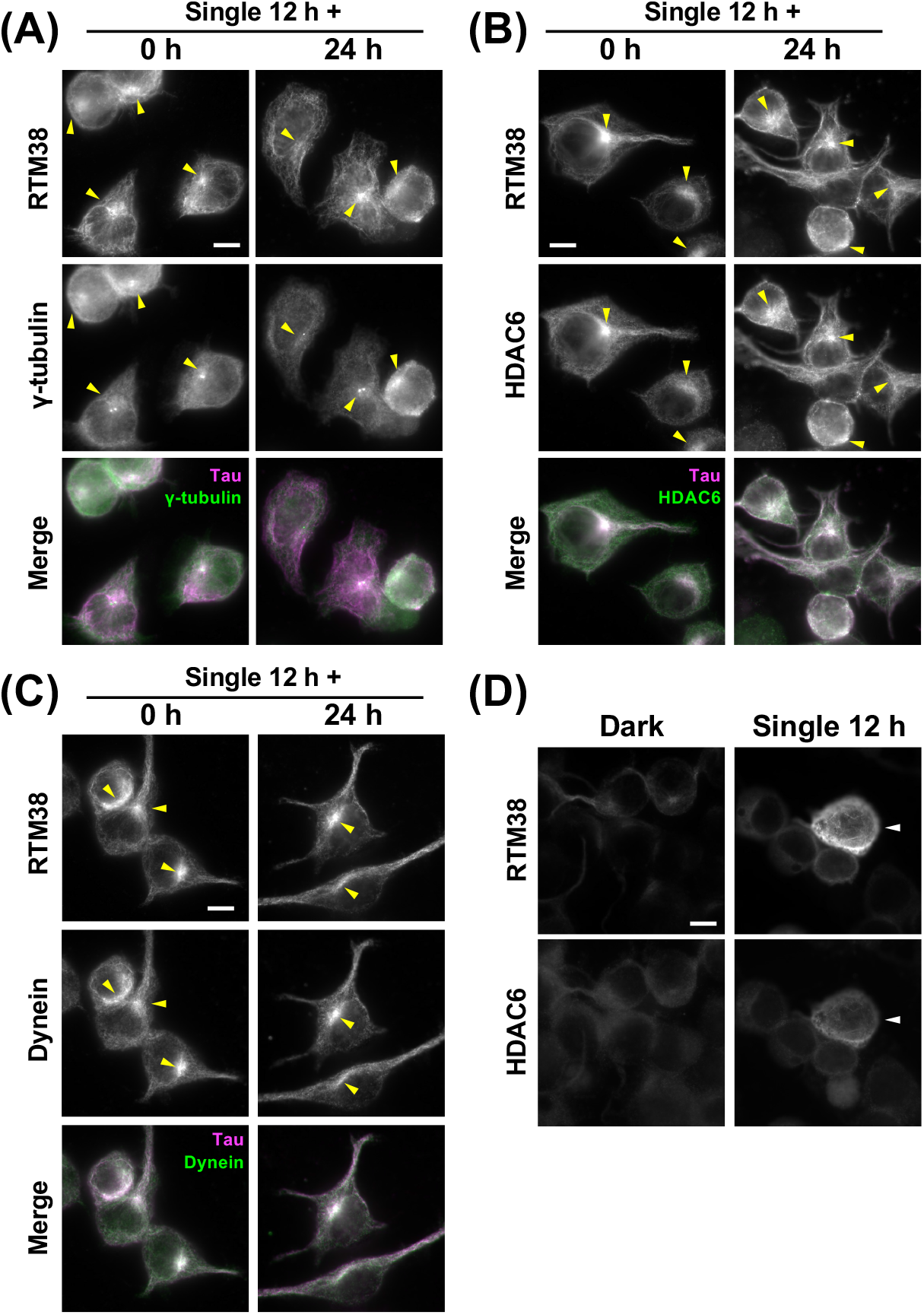
Co-localization of tau with aggresome proteins after blue light illumination termination. Multicolor immunofluorescence images of tau and aggresomal proteins in OptoTauKI cells immediately (0 h) and 24 h after terminating a single 12-h blue light illumination. Aggresomal proteins labeled were **(A)** γ-tubulin, (B) HDAC6, and **(C)** Dynein. Yellow arrowheads represent strong bright spots of tau. **(D)** Tau (RTM38, top) and HDAC6 (bottom) immunostaining under dark conditions (Dark, left) and immediately after a single 12-h light exposure (Light, right). The white arrowhead indicates a cell with a high expression level of tau and HDAC6. Scale Bars indicate 10 μm.

The process of transporting degenerated proteins along the microtube to the aggresome at MTOCs involves the microtubule motor protein dynein and histone deacetylase 6 (HDAC6), which recruits denatured proteins to dynein [34]. We examined the co-localization of these aggresome marker proteins with tau to investigate the possibility that the solid bright spots of tau that appeared in OptoTauKI cells after blue light arrest were contained in aggresome and its precursors. Tau signal co-localized with HDAC6 and dynein in the near-nuclear bright spots (arrowheads) and thick filaments (Figures 4B and 4C, 0 h). Tau exhibited stronger co-localization with HDAC6, or dynein, near the nucleus and in the cytoplasm 24 h after blue light termination. These results indicate that tau proteins are recruited to aggresome-associated proteins and ultimately accumulated to the aggresome in OptoTauKI cells with single 12-h continuous blue light exposure.

Western blot experiments demonstrated an increase in 150-kDa OptoTau after a single 12-h illumination (Figure 2D). RTM38 antibody immunostaining revealed cells with increased tau after a 12-h light exposure compared to dark conditions (Figure 4D, top). Interestingly, cells with increased tau upon light illumination demonstrated increased HDAC6 protein levels (Figure 4D, bottom). This result suggests that the increase in tau levels and filamentous tau induced by light stimulation may activate aggresome formation.

### Stable tau aggregation induced by intermittent blue light illumination

Continuous blue light illumination for 12 h (single 12 h) induced tau translocation to the aggresome, where denatured proteins accumulated but did not form stable aggregates, as seen in AD and other tauopathy. We then investigated conditions that would induce stable tau aggregation in OptoTauKI cells. Considering that 0.17–0.5-Hz repeated blue light stimulation induced stable aggregation of α-synuclein fused with CRY2 deliberative “CRY2Clust” [35], we expected that repeated cluster formation and dissociation of OptoTau would be effective in increasing the probability of tau molecules interacting in an optimal orientation to form oligomers through the microtubule-binding domain. Given that the 23.1-min half-life for CRY2olig cluster dissociation [21], we conducted the second illumination pattern, the “break and repeat,” composed of a 24-h illumination sequence consisting of a 16-h dark period (break) followed by eight 30-min dark/30-min blue light cycles (repeat), which was continued for 6–8 days (Figure 5A). OptoTauKI cells stably express the OptoTau protein, thereby enabling prolonged blue light illumination, which was impossible in transfected cells in previous studies.

**Figure 5.**
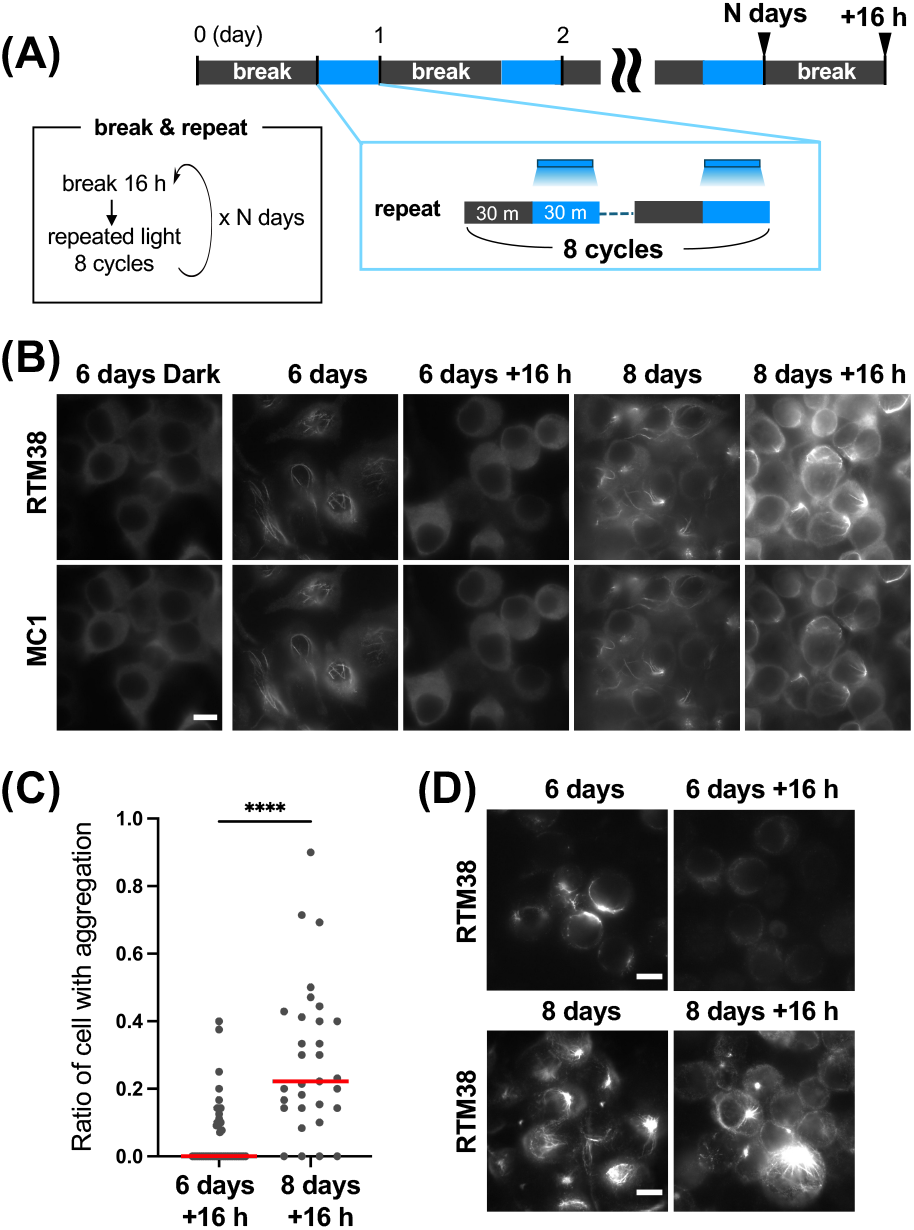
Formation of stable tau aggregates by break and repeat blue light illumination. **(A)** Blue-light illumination protocol defined as “break and repeat.” OptoTauKI cells were cultured in darkness for 16 h (break), followed by “repeat,” 8 cycles of 30-min darkness/30-min blue light illumination. The break and repeat protocol was conducted for N days. Arrows indicate the timing of fixation. **(B)** Immunofluorescence staining using total tau antibody (RTM38) and the antibody recognizing tau conformation change (MC1). Cells were fixed after being cultured in the dark for 6 days (6 days dark), at the end of 6 day’s repeat (6 days), after the 16-hour break on day 6 (6 day+16 h), at the end of 8 day’s repeat (8 days), after the break on day 8 (8 day +16 h). Bar indicates 10 μm. **(C)** Percentage of cells with stable tau aggregation 16 h after repeated blue light exposure. The red lines represent the median. 6 days +16 h: n = 46; 8 days +16 h: n = 31. ****: *p* < 0.0001, Mann–Whitney U-Test. **(D)** Methanol-resistant Tau at 6 days and 6 days +16 h (top), 8 days and 8 days +16 h (bottom), visualized by RTM38 antibody. Scale bar indicates 10 μm.

Total tau labeled with RTM38 antibody exhibited a thick linear filament structure immediately after the break and repeated illumination protocol for 6 days and 8 days (Figure 5B, RTM38). The stability of this filamentous structure of tau aggregation-like structures was evaluated based on their presence after a 16-h break following final repeat illumination (Figure 5A). Filamentous tau was not seen in OptoTauKI cells with a 6-day illumination followed by a 16-h break (6 days + 16 h) but remained in cells with an 8-day break and repeat plus 16 h break (8 days + 16 h) (Figure 5B). The MC1 antibody recognizes the initial pathological tau conformation [36,37]. The MC1 antibody labeled RTM38-positive filamentous structure immediately after terminating repeated illumination and after a 16-h illumination cycle for 8 days (Figure 5B, MC1). The percentage of cells with bright solid spots demonstrating stable tau aggregation after a 16-h break was significantly higher following 8 days of illumination than after 6 days (Figure 5C). The existence of stable tau in OptoTauKI cells after the break and repeat illumination protocol was evaluated by the methanol fixation, which removes soluble tau in the cytoplasm [27]. RTM38-positive aggregation in the cytoplasm was observed after 6 days of break and repeat, but this aggregation disappeared after a 16-h break (Figure 5D, Top). The amount of stable tau that remained after methanol fixation was increased immediately after 8 days compared to after 6 days of break and repeat illumination and was detected as thick filament structures (Figure 5D, 8 days). This filament-like methanol-resistant tau aggregation remained after a 16-h break (Figure 5D, 8 days + 16 h). The results indicate that break and repeated illumination over 8 days generated stable tau aggregation.

### Acceleration of stable tau aggregation formation by short light-dark cycles

Finally, we investigated the association between the temporal pattern of blue light/dark cycle repetitions and the time to induce stable tau aggregation. Here, 25-min dark/5-min blue light illumination cycles were repeated 48 times (48 repeats, Figure 6A). The total required time was 24 h, with a 4-h total illumination time. A total of 48 repeats of illumination protocol induced thick tau filaments as detected by RTM38 and MC1 antibodies in OptoTauKI cells (Figure 6B, left). Tau filaments generated by 48 repeats protocol remained after a 16-h break (Figure 6B, bottom). A total of 48 repeat illumination protocols resulted in the accumulation of CRY2olig-SNAP protein, control for OptoTau, into small spots, which were distinct from OptoTau filaments induced by the same illumination protocols, and these spots disappeared after a 16-h break (Supplemental Figure 3). This indicates that CRY2olig-SNAP alone could not induce stable aggregation formed by the 48 repeated illumination protocol.

**Figure 6.**
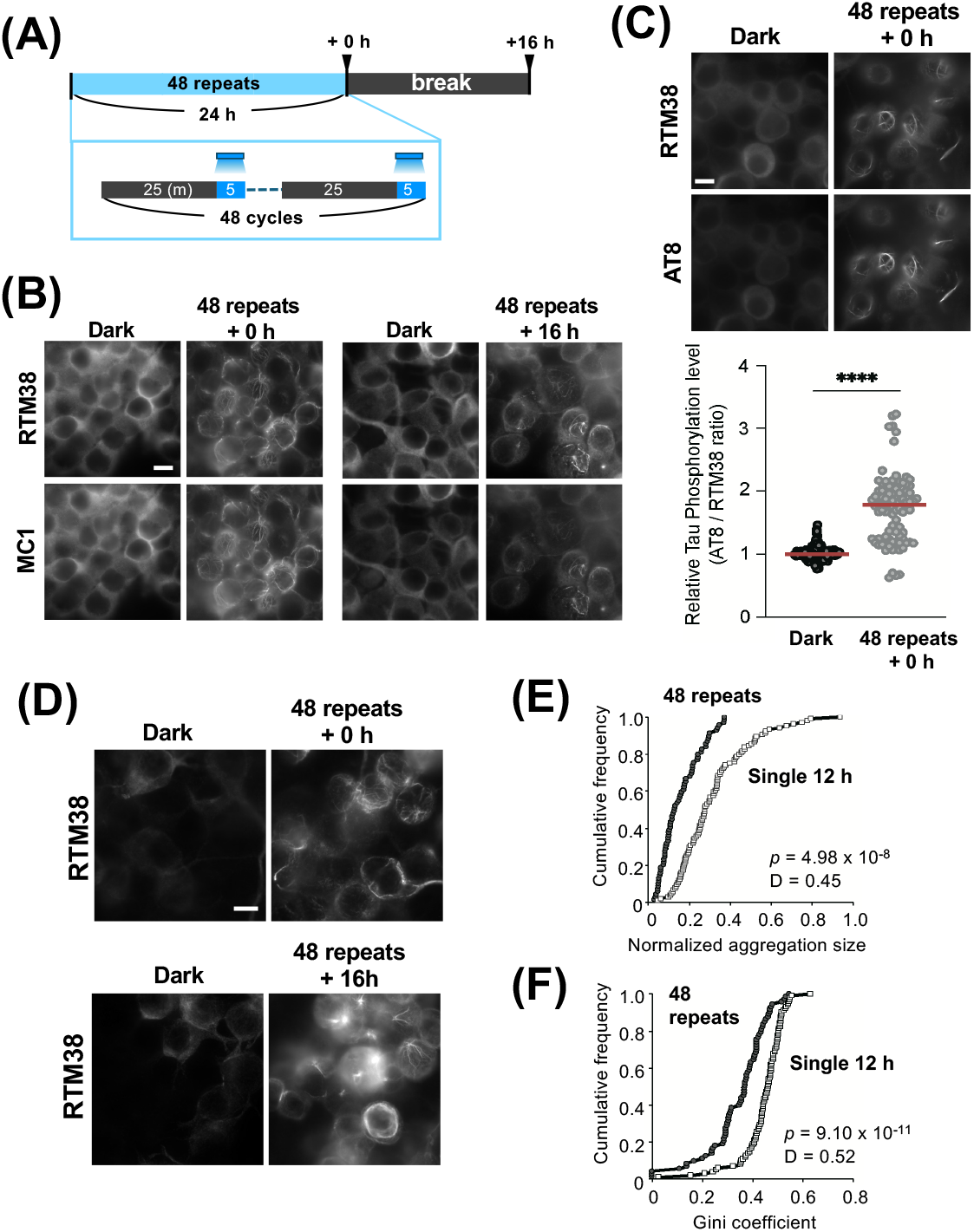
Acceleration of stable tau aggregation formation by repeated blue light illumination. **(A)** The 25-min dark and 5-min blue-light illumination cycle was repeated 48 times (48 repeats, 24 h). Cells were fixed at the timing indicated by arrowheads: at the end of 48 repeats (+0 h) or after 48 repeats, followed by a 16-h break (48 repeats +16 h). **(B)** Immunocytochemistry with RTM38 and MC1 antibodies of the cells fixed at the end of 48 repeats (48 repeats +0 h) and those fixed with PFA after a 16-h break (48 repeats +16 h). Cells cultured in the dark and fixed simultaneously (Dark) were utilized as the control. **(C)** Top: immunostaining with RTM38 and AT8 of the cells with 48 repeats (48 repeats +0 h) and control cells (Dark). Bottom: ratio of AT8 intensity to RTM intensity in the control cells (Dark, n = 184) and cells with repeated illumination (48 repeats, n = 91). The red line denotes the median for each group. ****: *p* < 0.0001, Mann–Whitney U-test. **(D)** Visualization of methanol-resistant tau by RTM38 antibody immediately after the end of repetitive illumination (48 repeats +0 h) or after 48 repeats followed by the 16-h break (48 repeats +16 h) and dark condition (Dark). Scale bars in **(B), (C)**, and **(D)** indicate 10 μm. **(E)** Cumulative plot of normalized tau aggregation size, tau aggregation width normalized to the nuclear axis (Supplemental Figure 1A) for cells with a 12 h-continuous illumination (12h single, open square, n = 104), and cells subjected to 48 repeats (48 repeats, filled circle, n = 69). p = 4.98 × 10^−8^, D = 0.45, Kolmogorov–Smirnov test. **(F)** Cumulative plot of normalized Gini coefficient (Supplemental Figure 1D), emphasizing the difference in the heterogeneity of the RTM38 signal intensity between cells with a 12-h illumination (Single 12 h, open square, n = 104) and cells with 48 repeats (48 repeats, filled circle, n = 73). p = 9.11 × 10^−11^, D = 0.52, Kolmogorov–Smirnov test.

Filaments labeled by the RTM38 antibody in cells with 48 repeats were also detected by the antibody recognizing phosphorylated tau (Figure 6C, top). The relative level of phosphorylated tau significantly increased after the 48-repeat illumination protocol (Figure 6C, bottom). The immunolabeling with RTM38 after methanol fixation verified the stability of filamentous tau in OptoTauKI cells immediately after the end of the cycle and after a 16-h break after 48 repeats (Figure 6D). These results indicate that stable tau aggregation could be induced in only 24 h by repeating the 25-min dark/5-min blue light exposure cycle.

In contrast to the thick tau accumulation in the aggresome near the nucleus induced by a continuous 12-h illumination (Figure 4), tau aggregates induced by 48 repeats of the 5-min light/25-min dark cycle were frequently located peripherally as thinner filaments (Figure 6C). The size of tau aggregates near the nucleus, calculated as indicated in Supplemental Figure 1A, was significantly larger in cells with a single 12-h illumination than in those with a 48-repeat illumination protocol (Figure 6E). We finally analyzed the Gini coefficient, which is an index of inequality originally utilized by social scientists but is now popularly used in the natural sciences, from genetics to astronomy [29], and obtained from the pixel fluorescence intensity of RTM38, to further investigate the effect of the temporal pattern of light illumination on the nature of tau aggregates in OptoTauKI cells (Supplemental Figure 1B, C). The cumulative plot of the Gini coefficient exhibited a Larger Gini coefficient in cells with a single 12-h illumination than in cells with 48 repeats (Figure 6F), indicating a significantly uneven distribution of tau fluorescence in cells with a single 12-h illumination, compared with the cell with 48 repeats. Altogether, these results indicate that the temporal pattern of blue light determines the characteristics of tau aggregates induced in OptoTauKI cells; 12-h single illumination induced the accumulation of tau in the aggresome. In contrast, 48 repeats formed stable tau aggregates in the broader area of the cell.

## Discussion

The present study generated cells knocking in OptoTau, which is an optogenetic tool based on the CRY2olig module, and developed a technique to manipulate tau aggregation with blue light. We discovered light illumination conditions that specifically induce tau accumulation in aggresomes or stable tau aggregation.

Recently, a series of studies revealed that chimeric protein transient overexpression combining tau with CRY2 or its derivative CRY2olig, i.e., families of OptoTau, causes the blue light-dependent formation of LLPS droplets and tau oligomers [23-26]. The present study manipulated tau by blue light in OptoTauKI cells, in which OptoTau was inserted into the genome by genome editing. The transfected plasmid is diluted by dividing cells, thereby attenuating the expression level of the product. Here, a cell line that stably expresses OptoTau enabled us to validate the blue light illumination protocol over several days while maintaining OptoTau expression.

A previous study has shown that tau condensate formed along microtubules within 50 s when SH SY5Y and HeLa cells overexpressing CRY2WT-mCherry-tau, a protein fusing CRY2 and human wild-type tau were exposed to blue light [23]. Consistent with this study, we found that tau formed filament structures colocalizing with microtubules in OptoTauKI cells exposed to blue light for 12 h. Conversely, a 3-min blue light exposure to primary cortical culture neurons overexpressing tau::CRY2, which is CRY2olig fused to the C-terminus of human wild-type tau, generated tau droplets and oligomers [24]. Such a tau droplet, which is considered an LLPS droplet, was induced by a 24-h blue light illumination in the presence of microtubule depolymerizing agent nocodazole in Neuro2a cells overexpressing our OptoTau, consisting of CRY2olig-Snap-human tau with P301L [25]. However, none of the light illumination protocols tested in the present study induced droplet-like structures in OptoTauKI cells. The differences in the OptoTau complex morphology induced by blue light may depend on multiple parameters, including the expression levels of OptoTau, cellular endogenous tau, blue light exposure time, exposure intensity, cell type, and microtubule integrity.

Part of the filamentous tau induced by 12-h blue light illumination on OptoTauKI cells accumulated in MTOCs stained with γ-tubulin and co-localized with aggresomal component proteins HDAC6 and dynein, as was observed in Neuro2a cells overexpressing OptoTau [25]. The increased HDAC6 amount in cells with elevated tau implies that selective autophagy is triggered as a stress response. Our study did not confirm tau ubiquitination up-regulation, which is a signal for transport to aggresome. Additionally, increased aggresome activity following light stimulation, which causes OptoTau and endogenous tau degradation, remains unclear. Previous studies demonstrated increased phosphorylated tau and tau oligomers in the hippocampus of mice in which the aggresomal protein p62 is knocked out [38]. Optogenetically accumulated OptoTau co-localizes with endogenous p62 [25]. The role of p62 in light-induced aggresome formation and tau degradation warrants further investigation.

This study indicated two illumination conditions that induce methanol-resistant tau aggregation: one is to repeat a 24-h illumination sequence consisting of a 16-h dark period followed by eight 30-min dark/30-min blue light cycles (break and repeat) for 8 days, and another is repeating 25-min dark/5-min blue light cycles for 24 h (48 repeats). This aggregation did not disappear even after a 16-h light illumination termination. Therefore, these tau aggregates are interpreted as a stable residual structure. Stable filamentous tau aggregates were observed by the antibody recognizing the initial conformational change in pathological tau (MC1) and the antibody detecting pathological phosphorylation of tau (AT8), indicating that these aggregates include pathological tau. Previously we found that OptoTau with the truncated N-terminal of tau optogenetically induced sarkosyl-insoluble tau aggregation, whereas OptoTau with full-length tau did not cause insoluble tau aggregation [25]. The N-terminal of tau inhibits full-length tau polymerization [39]. Thus, OptoTau which lacks its N-terminal is more likely to produce aggregation seed. Whether methanol-insoluble stable tau aggregates induced by repeated blue light illumination in the present study were sarkosyl-insoluble remained unclear. However, the finding that methanol-resistant tau persisted for at least 16 h after light cessation indicates that OptoTau could function as a new tool for inducing stable aggregates from full-length tau at the indicated time.

The present study demonstrated that differences in the temporal pattern of light illumination produce variations in the spatial pattern of OptoTau aggregation. Previous research revealed that repeated light stimulation at 0.17 Hz induced stable aggregates of opto-α-synuclein, which is α-synuclein fused with CRY2clust, in human iPS cell-derived neurons [35]. Taken together with the results of this study, which exhibited that only discontinuous repeated light illumination rather than continuous light illumination caused stable OptoTau aggregation, repeated light stimulation with interruptions not only reduces phototoxicity but also promotes the aggregation effect of CRY2 fusion proteins. The frequent OptoTau concentration by blue light through CRY2olig, followed by dissociation during the dark condition, may increase the possibility of interaction between tau molecules at appropriate positions to form β-sheet structure through its microtubule-binding domain, which is located in the core of tau filaments [40]. The temporal pattern, including the duration of one illumination, interval time, and number of repetitions, is considered another crucial factor in introducing stable aggregation with OptoTau. The 48 repeat protocol in 24 h induced stable aggregation, despite the shorter total time of blue light illumination than the break and repeat protocol for 8 days; 4 h for 48 repeats vs. 24 h for break and repeat. This result implies that tau aggregation can be induced more efficiently when CRY2olig binding/dissociation is rapidly repeated. However, 144 cycles of 1-min blue light/9-min dark illumination failed to cause fibrous tau aggregation, despite the high number of cycles (data not shown). This study only presented two conditions under which stable tau aggregation induction is possible, and the optimal aggregation induction system should be further explored.

This study showed that OptoTau exhibited different behaviors, such as aggresome formations and stable tau aggregation, i.e., proteolysis and protein aggregation, depending on the temporal pattern of the light stimulation. OptoTau presents a new experimental possibility for future introduction into the mouse brain, where the assembly state of tau can be manipulated with light at the intended timing in specific cells at the targeted brain region. Additionally, this optogenetic tool helped validate the existence of “singularity cells,” i.e., a small number of rare cells that cause significant changes throughout the system [18]. The neurons that first form tau aggregates in LCs and ECs may be considered “singularity cells” in AD [20]. OptoTau that specifically induces tau aggregation only in LCs and ECs will verify whether they can be “singularity cells” that spread tau pathology to the cortex.

There are some issues that need to be resolved before utilizing OptoTau as a model for tauopathies in the future. One is the tau isoform. The tau isoform, with the 31 amino acid repeats encoded by exon 10, is named the 4R isoform, and the isoform without the second repeat is called the 3R. The tau aggregates of AD and CTE contain 3R and 4R tau, whereas Pick’s disease is composed of only 3R tau, and CBD and PSP consist of only 4R tau. The molecular structure of the filaments within tau aggregates varies in each tauopathic disease [10]. The molecular structure of the tau aggregation core determines the tau aggregate properties (including the 3R tau to 4R tau ratio) specific to tauopathies [41]. Current OptoTau includes only 4R-type tau. In the future, developing 3R-type OptoTau will be necessary to establish optogenetic models specific to AD, CTE, and Pick’s disease. The 3R to 4R tau expression ratio in OptoTau-induced cells should be controlled. Additionally, optogenetically inducing aggregation nuclei specific to each tauopathy will be a significant challenge.

## Conclusion

This study used the optogenetic tool OptoTau, based on the CRY2olig module, to form tau aggregation in cells. Specific light exposure conditions introduced aggresome and stable aggregation in OptoTauKI cells. The present results provide a new technological basis for establishing a singular point of tau aggregation at a targeted time in specific cells.

## Conflict of interest

The authors declare no conflict of interest related to this publication.

## Author contributions

SS, TU, NK, TT, BX, AH, and YS performed the experiment. SY, AT, HY, GM, and HB were involved in the design and supervision of this work. SS, TU, NK, BX, AH, and HB processed the experimental data, performed the analysis, drafted the manuscript, and designed the figures. All authors discussed the results and commented on the manuscript.

## Data availability

The evidence data generated and/or analyzed during the current study are available from the corresponding author upon reasonable request.

A preliminary version of this work, DOI: https://doi.org/10.1101/2024.05.07.592868, was deposited in the bioRxiv on May 8th, 2024.

## Acknowledgments

The authors thank Prof. Peter Davies (Albert Einstein College of Medicine) for providing the MC1 antibody, Dr Sumiyo Mimura (AS ONE Corporation) for technical support of single cell cloning, and Dr. Naruhiko Sahara (National Institutes for Quantum Science and Technology) for insightful discussion. This research was funded by Grant-in-Aid for Scientific Research on Innovative Areas “Singularity Biology” (JP18H05410 to HY, JP18H05414 to HB) from the Ministry of Education, Culture, Sports, Science and Technology, Japan, JSPS Grants-in-Aid for Scientific Research JP20K06896, JP23K05993 to YS, JP21H02450, 23K18116 to HB, and grants from Naito foundation and Takeda Science Foundation to HB. DeepL, ChatGPT 3.5, Grammarly, and Enago the editing brand of Crimson Interactive Pvt. Ltd supported the English editing of this article.

## Notes

https://drive.google.com/file/d/1QKJZNkgaSatt4B2qsOOepWXK-Y2o34fa/view?usp=drive_link

https://drive.google.com/file/d/1_6KHoZsGIttftr7kqVhN9EOEUUtW61AT/view?usp=drive_link

### Competing Interest Statement

The authors have declared no competing interest.

### Summary of Updates

(1)The title was modified according to the reviewer's suggestion (2)We have defined the names of the three illumination protocols to clearly present the blue-light illumination conditions used in this study. (3)We added data on biochemistry and immunocytochemistry using the AT8 antibody to show the existence of pathological tau (Figure 2E and Figure 6C). (4)We performed Western blot analysis under reductive conditions and added the data with another pan-tau antibody (Figure 2C, 2D and Supplemental Figure 2). (5)Immunocytochemistry data were replaced with datasets in which the cell density of the control and blue-light-illuminated groups were equivalent (Figure 2A). (6)Novel quantification indexes were induced to analyze the difference in tau aggregation induced by different illumination protocols (Figure 6E, 6F, and Supplemental Figure 3). (7)In line with the above changes, we modified the main text, corrected the typing errors, and added new references. (8)Due to the additional experiment, we included one more author (T. T.) who carried out a biochemical analysis.

